# Matrix mechanics, not hypoxia, modulate quiescin sulfhydryl oxidase 1 (QSOX1) in pancreatic tumor cells

**DOI:** 10.1101/2022.10.19.512796

**Authors:** Catherine S. Millar-Haskell, Colin Thorpe, Jason P. Gleghorn

## Abstract

Pancreatic ductal adenocarcinoma (PDAC) is the 4th leading cause of cancer-related deaths in the U.S., despite only being the 11th most common cancer. The high mortality rates of PDAC can be partially attributed to the tumor microenvironment. Unlike most carcinomas, PDAC is characterized by a strong desmoplastic reaction, or a fibrotic stiffening of the extracellular matrix (ECM) in response to chronic inflammation. The desmoplastic reaction is mediated by cancer-associated fibroblasts that deposit ECM proteins (collagens, laminins, fibronectin, etc.) and secrete matrix-remodeling proteins in the tumor parenchyma. Within the past decade, the enzyme quiescin sulfhydryl oxidase 1 (QSOX1) has gained recognition as a significant contributor to solid tumor pathogenesis, but its biological role remains uncertain. QSOX1 is a disulfide bond-generating catalyst that participates in oxidative protein folding in the mammalian cell. Current studies show that inhibiting or knocking down QSOX1 reduces pancreatic cancer cell migration and invasion, alters ECM deposition and organization, and decreases overall tumor growth in mice. However, it is unclear which features of the tumor microenvironment modulate QSOX1 and cause its overexpression in cancer. In this study, we explored potential regulators of QSOX1 expression and secretion by testing two major features of PDAC: hypoxia and mechanical stiffness. To induce hypoxia, we exposed pancreatic cancer cells to atmospheric (low O_2_) and chemical (CoCl_2_) hypoxia for up to 48 hours. QSOX1 gene and protein expression did not change in response to hypoxia. Substratum stiffness was modulated using polyacrylamide gels to represent the dynamic pathological range of elastic moduli found in PDAC tissue. We discovered that QSOX1 levels were decreased on softer surfaces compared to conventional tissue culture plastic. This paper presents new results and challenges prior findings on QSOX1 regulation in pancreatic tumor cells.

## INTRODUCTION

PDAC is a lethal cancer that typically develops from pancreatic lesions in chronic pancreatitis^1,2^. PDAC is currently the third leading cause of cancer-related deaths in the United States and is predicted to surpass colon cancer by 2030^3,4^. One of the most defining features of PDAC is a desmoplastic reaction that resembles an intense fibrotic wound healing response^5–7^. During desmoplasia, activated fibroblasts proliferate and deposit excessive ECM proteins to form a dense stroma around the tumor site. The ECM proteins that comprise the pancreatic stroma include collagens I, III and IV, laminin, and fibronectin and are often disorganized. The stroma in PDAC creates a physical barrier that results in poor vascularization, hypoxia, and increased mechanical stiffness^8,9^. This ultimately impairs tumor perfusion, delivery of anti-tumor drugs, and effectiveness of chemoradiotherapy. Fundamentally, the desmoplastic response is a key driving force behind the lethality of PDAC.

Matrix remodeling enzymes are significantly involved in the tumor microenvironment^10^. Matrix metalloproteinases (MMPs), which catalyze the degradation of ECM proteins, are implicated in local cell invasion and fibroblast activation^11–15^. There are over 17 different MMPs, and at least nine are involved with pancreatic cancer^12,16^. Lysyl oxidases (LOXs) are a class of copper-dependent enzymes that catalyze collagen and elastin crosslinking, which increases the local tensile strength of tissue. Abnormal LOX expression is strongly implicated in fibrotic disease progression in cancer^17–21^. QSOX1 is an enzyme that catalyzes the generation of disulfide bonds and is overexpressed in multiple solid tumors^22^. Notably, QSOX1 expression is associated with cancer invasion^23–25^ and metastasis^26–29^. There is also evidence that QSOX1 participates in matrix remodeling events by altering ECM protein deposition^30,31^, although this mechanism has not been fully elucidated. Additionally, while there has been significant progress toward understanding the role of QSOX1 in solid tumors, its presence in the stroma remains somewhat of a mystery. Specifically, it is not well understood why QSOX1 is overexpressed in solid tumors, what features of the tumor microenvironment modulate its expression, and how secreted QSOX1 functions in the extracellular space.

One of the hallmarks of desmoplasia is a mechanical stiffening of the tumor. The elastic modulus of human pancreatic tissue falls between 0.5-1.5 kPa, while PDAC tumors range from 1.5-13.0 kPa^32^. The elastic moduli of healthy murine pancreata were recorded at 1 kPa with a modest increase of roughly 4 kPa in cancerous tissue^33^. In humans, the stiffness of PDAC tissue varies both locally within a specimen and across patients but appears to be 3-30 times stiffer than healthy pancreatic tissue. Cell populations in the PDAC tumor microenvironment are mechanically responsive. In one study, pancreatic stellate cells cultured on 1 kPa polyacrylamide (PAA) gels maintained a quiescent state, while those cultured on 25 kPa PAA gels adopted a fibroblastic phenotype^34^. Pancreatic cancer cells cultured on PAA gels (1 kPa, 4 kPa, and 25 kPa) likewise responded differentially to matrix stiffness^33^. Cells on softer gels were found to significantly downregulate connective tissue growth factor expression, which is involved in hypoxia-mediated tumor fibrosis. In another study, pancreatic cancer cells cultured on crosslinked collagen gels increased membrane-tethered MMP activity in a contractility-dependent manner, and cell spreading was correlated with higher MMP activity^35^. While QSOX1 has been linked with cell adhesion, invasion, and high-grade tumors, no studies have investigated whether matrix mechanics directly regulates QSOX1 protein expression and secretion.

A second hallmark of PDAC is hypoxia, or low tissue oxygenation. In 2000, a small clinical study found that the median oxygen concentration of resected pancreatic tumors was below 0.6% (3 mm Hg) compared to 1-12% in adjacent tissue^36^. Hypoxia inducible factor 1 (HIF-1) is a transcription factor that contains two subunits, HIF-1α and HIF-1β. The β subunit is constitutively expressed, while the α subunit contains an oxygen-dependent degradation domain that signals it for proteasomal degradation under normoxic conditions. During hypoxic conditions, HIF-1α accumulates in the cytosol and translocates into the nucleus where it dimerizes with HIF-1β and promotes transcription of the target gene. HIF-1α reportedly has numerous target genes, including vascular endothelial growth factor^37^ (angiogenesis), sonic hedgehog^38,39^ (fibrosis), LOX^40^ and MMPs^41^ (matrix remodeling), fascin^42^ (invasion), and connective tissue growth factor (fibrosis). Many of these genes have the putative hypoxia-response element (HRE), a sequence that contains 5’-TACGTG-3’ and is recognized by HIF-1^43^. Interestingly, the QSOX1 gene in humans has multiple HREs (promoter sequence 5’-RCGTG-3’). In 2013, QSOX1 was reported to be upregulated in hypoxic conditions in pancreatic cancer cells in a HIF-1α-dependent manner^44^. However, additional studies investigating hypoxia-mediated QSOX1 upregulation have not been conducted.

In this study, we investigated QSOX1 protein and gene expression resulting from changes in matrix stiffness and hypoxia in pancreatic cancer (PANC-1) cells. To test the effect of matrix stiffness, we synthesized PAA gels with a broad range of elastic moduli representing the pathological range of human PDAC (2-60 kPa) and compared it against traditional tissue culture plastic and cover glass. To mimic hypoxia, we used both atmospheric (1% O2) and chemical (400 μM CoCl2) induction and measured changes to QSOX1 gene expression and intracellular protein levels. Surprisingly, QSOX1 expression was not responsive to hypoxic challenges. However, we did see significant downregulation of QSOX1 on softer PAA gels compared to stiff surfaces, with a corresponding decrease in total secreted QSOX1. We thus present the novel finding that matrix mechanics may regulate QSOX1, but not necessarily hypoxia, in pancreatic tumor cells. This work provides an alternate explanation for why QSOX1 is overexpressed in PDAC and contextualizes its observed effects on cell-ECM interactions.

## MATERIALS AND METHODS

### Cell culture

Pancreatic cancer (PANC-1, ATCC) cells were cultured in Dulbecco’s Modified Eagle Medium (DMEM, Sigma-Aldrich) supplemented with 10% FBS (Atlanta Biologicals) and 1% penicillin/streptomycin (“DMEM”). This media formulation was used for hypoxic studies. A low-serum modified cell line was previously established where PANC-1 cells were cultured in DMEM supplemented with 2% FBS and 1% Insulin-Transferrin-Selenium Media Supplement (ITS, R&D Systems) (“DMEM-ITS”)^45^. This media formulation was used for experiments that used conditioned media from cells cultured on PAA gels to reduce biofouling and minimize serum shock during serum starvation in experimental conditions. Cells were maintained in a humidified chamber at 37°C and 5% CO_2_ during routine culture.

### Synthesis of PAA gels

Surface functionalization and PAA polymerization were adapted from previously established methods^46–48^. The process for creating the PAA gels is summarized in **Figure 1**. In this study, two PAA gel formats were used: a 22×22 mm coverslip for intracellular protein quantitation and a large dish for collecting conditioned media. PANC-1 cells were seeded on ECM-coated PAA gels or directly on ECM-coated coverslips (20k cells/cm^2^) in DMEM or 145 mm dishes (10k cells/cm^2^) in DMEM-ITS.

**Figure 1:**
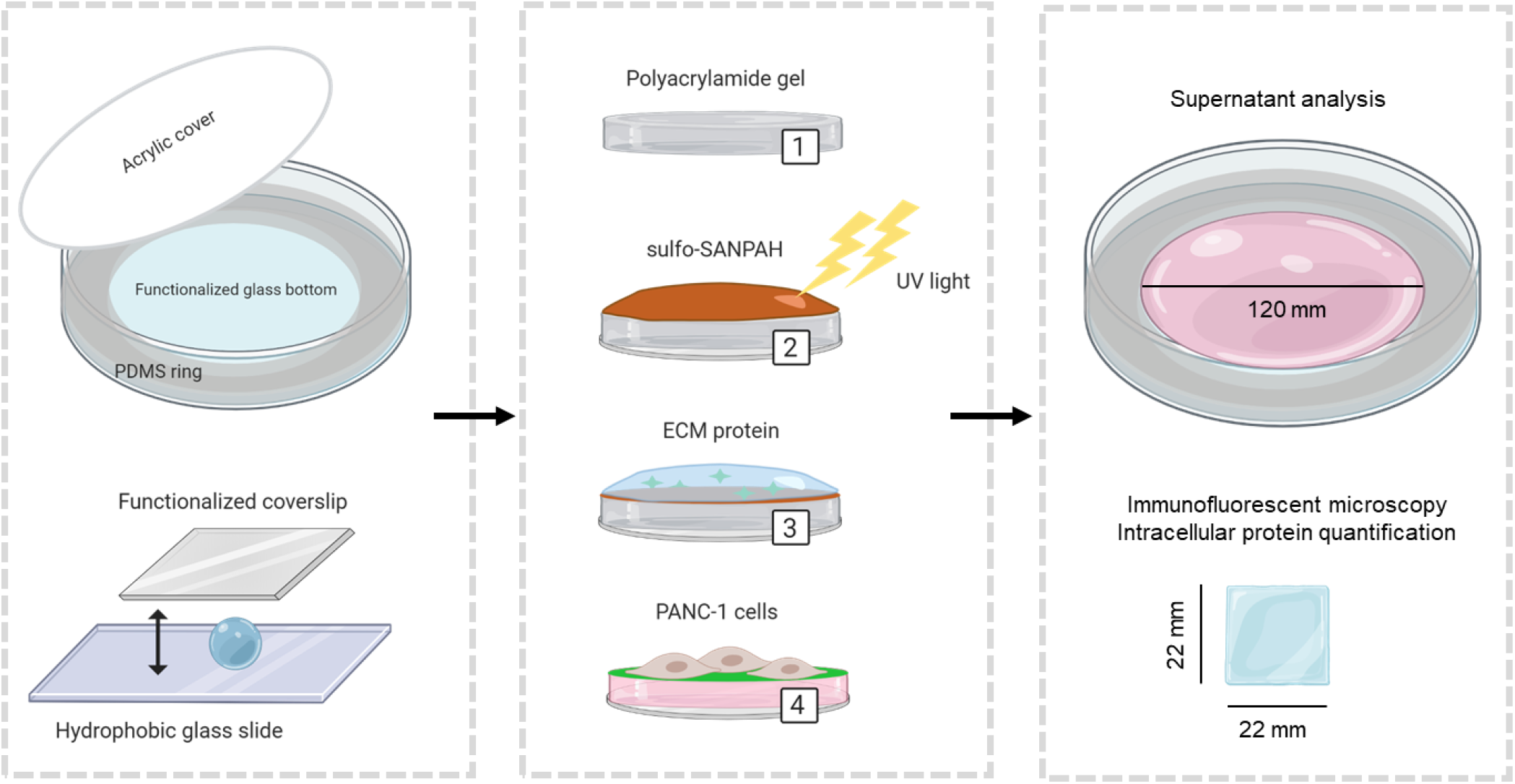
Process for synthesizing PAA gels. Left panel: for the scaled-up gel platform, a sterile PDMS ring was fitted into a functionalized glass petri dish and PAA solution was sandwiched between the glass and acrylic cover. For the small platform, PAA solution was sandwiched between small 22×22 mm functionalized coverslips and a hydrophobic glass slide. Middle panel: (1) PAA was polymerized for 20 minutes, and the acrylic cover or coverslip was removed. (2) The crosslinker sulfo-SANPAH was activated with 365 nm light for 10 minutes. (3) The ECM solution of choice was incubated overnight at 37°C. (4) Cells were seeded on top and incubated for 2 h to facilitate attachment before adding DMEM-ITS media. Right panel: the coverslips were used for immunofluorescence and intracellular QSOX1 quantitation while PAA dishes were used for conditioned media production and supernatant analysis. Created with Biorender.com

### Creation of polydimethylsiloxane (PDMS) ring molds (Large format PAA gels)

To create PDMS molds, 27 g of Sylgard 184 elastomer base and 3 g of Sylgard 184 curing agent were mixed and poured into a 145 mm petri dish^49^. The mixture was de-gassed under vacuum for 30 min to remove air bubbles and cured overnight at 65°C. The cured PDMS measured ~1.5mm thick using a digital caliper. The PDMS was cut into rings with an inner diameter of 120 mm, cleaned sequentially with isopropanol and ethanol, and autoclaved with a 30 min sterilization cycle.

### Surface functionalization

Large 145 mm glass petri dishes (VWR; 25354-127) or 22×22 mm coverslips (VWR; 16004-302) were functionalized to facilitate attachment of PAA to surfaces. Coverslips were functionalized in bulk using a staining rack (Thermo Fisher), large rectangular glass dishes, and a stir bar according to Aratyn-Schaus et al.^47^. Glass petri dishes were functionalized by adding solutions directly into the dish. The proceeding steps were identical for both types of platforms. First, surfaces were immersed in 0.1 M NaOH for 30 min, and after removal of the solution, were allowed to air dry. Next, surfaces were submerged in 2% 3-aminopropyltriethoxysilane (APTES) in isopropanol for 10 min. To fully remove unreacted APTES, surfaces were soaked in ddH_2_O for 40 min with four total exchanges of ddH_2_O (10 min each). Surfaces were immersed in 0.5% glutaraldehyde in ddH_2_O for 30 min and then allowed to fully dry prior to PAA synthesis.

### Polymerization of PAA gels

Two platforms were used: a 22×22 mm coverslip and a scaled-up version utilizing a reusable 145 mm glass dish. Pre-polymer solutions of 40% acrylamide (Sigma Aldrich; A4058) and 2% bis-acrylamide (Sigma Aldrich; M1533) were prepared according to the predicted storage moduli. Solutions were degassed in a vacuum chamber for 30 min to lower the concentration of dissolved oxygen. A 10% stock solution of ammonium persulfate (APS) in ddH2O was prepared in advanced and stored at −20°C for up to 6 months. To initiate polymerization, PAA polymer solutions were removed from the vacuum chamber and the co-initiators APS (0.1% final concentration) and tetramethylethylenediamine (TEMED; 0.01% final concentration) were added. For coverslips, solutions were mixed and then quickly pipetted (50 μl) onto Rain-X®modified glass slides. The functionalized coverslips were placed on top of these PAA solutions and allowed to polymerize for 20 min. For the 145 mm glass petri dishes, the sterile PDMS molds were inserted to act as a gasket and the PAA solution was added directly onto the functionalized glass dish. A laser-cut acrylic cover (125 mm diameter) was gently placed on top of the PDMS ring to create an airtight seal during polymerization. To remove the acrylic cover from fully polymerized gels, the unpolymerized solution at the PDMS-acrylic interface was vacuum aspirated and 100% isopropanol was added to fill the void and significantly reduce surface tension. Acrylic covers detached easily after successful infiltration of isopropanol between the PAA and cover. To remove 22×22 mm coverslips from the glass slide, the assembly was immersed in ddH2O and carefully pried off using forceps.

### Rheology

PAA solutions were polymerized directly on a DHR3 rheometer (TA Instruments) using a 20 mm parallel plate geometry and a PDMS mold (1.5 mm thick, 20 mm diameter well) that was affixed around the 20 mm upper plate. Measurements were taken using a 1 rad/s oscillatory frequency, 0.1% constant strain, and a 30 min time sweep. Young’s modulus (elastic modulus) was calculated from:

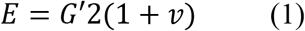

where E is Young’s modulus, G’ is the storage modulus, and *ν* is the Poisson’s ratio for the PAA gels (approximately 0.5) as reported in literature^48^.

### Coating ECM proteins on plastic surfaces

Fibronectin purified from bovine plasma (Pel-Freez Biologicals 37141-1), collagen (Corning, 354236), gelatin (Sigma-Aldrich G9391), laminin (Corning, 354232), and Matrigel (Corning, 354230) were coated on cell culture dishes as previously described^45^.

### ECM protein-PAA conjugation

ECM proteins were covalently attached to PAA using the crosslinker Sulfo-SANPAH (Covachem; 13414-100). To begin, the ECM protein of choice (fibronectin, gelatin, Matrigel, or collagen; 100 μg/ml) was diluted in HEPES-buffered saline (HBS; 50 mM HEPES with 0.9% NaCl, pH 8.0). Stock solutions of sulfo-SANPAH (10 mg/ml stored at −80°C) were diluted to 0.5 mg/ml in ddH_2_O and allowed to activate with the surface of the PAA gels under a 365 nm 100-Watt UV lamp (UVP) for 10 min. PAA gels with activated sulfo-SANPAH were washed twice with HBS. The gels were incubated in ECM solution overnight at 37°C in a cell culture incubator. After successful conjugation, the ECM solution was removed and replaced with HBS. Gels were soaked in HBS at 37°C overnight with a minimum of two exchanges to remove excess unconjugated sulfo-SANPAH and/or ECM protein and was visually inspected for a color change of burgundy to a light transparent orange.

### Fibronectin-TRITC (FN-TRITC) conjugation

The fluorescent dye 5(6)-tetramethylrhodamine isothiocyanate (TRITC; Alfa Aesar) was conjugated to fibronectin with slight modifications to the Thermo Fisher dye conjugation guideline. Briefly, 1 mg fibronectin in 0.1 M sodium bicarbonate buffer was reacted with 1 mg TRITC dye for 1 h at RT. The resulting FN-TRITC was purified with a PD-10 desalting column (Cytiva) equilibrated with PBS. Fractions were collected and analyzed for fluorescent intensity (Ex/Em 543/71) and protein concentration (OD 280). FN-TRITC was conjugated to PAA using the crosslinker sulfo-SANPAH. As a control, 20 μg/ml FN-TRITC was passively adsorbed to the glass. Successful conjugation to PAA was visualized using a Zeiss Axio Observer Z1 inverted epifluorescence microscope and quantified for fluorescent intensity using the histogram feature in ImageJ. The mean fluorescence of each coverslip was subtracted from the background fluorescence.

### Enzyme-linked Immunosorbent Assay (ELISA)

Cells on PAA gel coverslips were lysed with RIPA buffer (Alfa Aesar). For the ELISA, recombinant hsQSOX1 standard or 1 μg of lysate was diluted into coating buffer (50 mM sodium carbonate/bicarbonate, pH 9.6) and incubated overnight at 4°C in a sealed 96-well plate. Next, the plate was washed three times in wash buffer (PBS with 0.05% Tween-20). Samples were incubated with blocking buffer (PBS with 1% BSA) at RT for 1 h. The blocking buffer was removed, and samples were incubated with QSOX1 antibody (1:1000, ProteinTech) overnight at 4°C. The next day, the plate was washed four times in wash buffer and samples were incubated with anti-rabbit HRP-conjugated secondary (1:50,000, LI-COR) for 1 h. The plate was washed three times and developed with TMB Core (Thermo Fisher) for 30 minutes. The reaction was stopped with 0.2 M sulfuric acid and read with an H1 Synergy microplate reader at 450 nm.

### Generation of conditioned media

PANC-1 cells were sub-cultured in ECM-coated T175 culture dishes (Thermo Fisher) or collagen-PAA petri dishes in DMEM-ITS and maintained in a humidified chamber at 37°C and 5% CO_2_ for 24 h. To begin conditioned media production, the T175 flasks were washed twice with phosphate buffered saline (PBS). The PAA dishes were soaked with Hank’s Balanced Saline Solution containing calcium and magnesium (HBSS++, Corning) for 2 hours with 4 exchanges every 30 min to remove protein from the gel. To both types of dishes, 30 ml basal DMEM was added. After 24 h, the conditioned media was collected in 50 ml conical tubes and centrifuged at 300 x g for 10 min at 4°C to pellet detached cells. The conditioned media supernatant was transferred to new conical tubes and centrifuged 2000 x g for 20 min to remove large cellular debris. This step was repeated at least three times with decantation to new conical tubes prior to each spin. The clarified conditioned media was dosed with 2 μl 100x HALT protease inhibitor cocktail (Thermo Fisher), diluted with an equal volume of HBS (30 ml), and concentrated to 200 μl using a 10kDa MWCO spin filter concentrator (Cytiva, 28932225). Cells that were attached to the culture dishes were trypsinized and counted on a NovoCyte flow cytometer (Agilent).

### Nanoparticle Tracking Analysis (NTA)

Sample size distributions and concentrations were measured via nanoparticle tracking analysis (NTA; NanoSight NS300; Malvern, UK). NTA was used to check for consistency of size distributions and concentrations among the batch isolations. Samples were diluted in sterile cell culture grade PBS and processed using the following parameters: three 45-s videos were recorded per sample with a camera level of 15. Software settings were kept consistent between sample batches (screen gain 1.0, threshold 8) (Nanosight NTA 3.3).

### DC protein assay

The protein concentration for all samples was quantified using the DC protein assay (Bio-Rad). Briefly, 5 μl of BSA standard (0.2 – 2 mg/ml) or sample was pipetted into a 96-well plate along with 25 μl reagent A’ and 200 μl reagent B. Samples were read after 15 minutes using an H1 Synergy microplate reader (Gen5 3.11) with absorbance of 562 nm.

### Hypoxia studies

PANC-1 cells were seeded in tissue culture 6-well plates (200k cells/well) in DMEM. Hypoxia studies were conducted 24 h post-seeding. For chemical induction of hypoxia, PANC-1 cells received 400 μM CoCl_2_ diluted in media for up to 48 h. For atmospheric induction of hypoxia, PANC-1 cells were placed in a humidified hypoxic chamber at 37°C, 5% CO_2_, 1% O_2_, and 94% N_2_.

### Isolation of mRNA and real-time reverse transcription quantitative polymerase chain reaction (RT-qPCR)

The mRNA from hypoxia studies were isolated using TRIzol™ reagent and purified using a Bioline mRNA purification kit. Samples were processed via RT-qPCR using QSOX1 (Hs00170210_m1) and RPLP0 (Mm00725448) TaqMan probes (Thermo Fisher) and Bioline amplification mix. The ΔΔCt values were calculated from first normalizing to internal gene expression (RPLP0) and then normalizing to the experimental control of interest (time or normoxia). Fold change (2 ^-ΔΔCt^) was calculated.

### Western blotting

SDS-PAGE was performed using the Quadra Mini Vertical Blotting System (Expedeon, Abcam) according to the manufacturer’s instructions. All western blotting reagents were purchased from Expedeon unless otherwise noted. Samples were mixed with 4x lithium dodecyl sulfate (LDS) sample buffer, 1.0% Triton X-100, and 15 mM dithiothreitol and heated at 95°C for 10 min. Samples were loaded either by equal protein (3.5 μg) or equal particle count into each lane of a RunBlue 4–12% TEO-Tricine Protein Gel and run at 130 V in RunBlue TEO-Tricine run buffer for 1.5 h with a stir bar and ice pack. The proteins on the gel were transferred to a 0.2 μm pore size nitrocellulose membrane (GE Healthcare) for 10 min using a Power Blotter system (Invitrogen, Thermo Fisher Scientific). Membranes were blocked with 5% Omniblok™ dry milk (AmericanBIO) in tris buffered saline (TBS) with 0.1% Tween-20 (TBS-T) overnight at 4°C with gentle rocking. After blocking, the membrane was incubated overnight at 4°C or with primary antibody solution (1:1000 dilution for anti-Alix [E4T7U, Cell Signaling Technology], anti-QSOX1 [12713-1-AP, ProteinTech], anti-HIF-1α [610958, BD Biosciences], β-tubulin [Cell Signaling, 2128S], and anti-ITGB1 [P5D2, Abcam], 1:2000 dilution for anti-GAPDH [60004-1-Ig, ProteinTech], and 1:5000 dilution for β-actin [8H10D10, Cell Signaling Technology]). The membranes were washed three times with TBS-T for 5 min and incubated with the corresponding HRP-conjugated secondary antibody (anti-mouse or anti-rabbit, LI-COR) for 1 h at RT (1:100000). Membranes were washed thrice with TBS-T for 5 min each and twice with TBS for 5 min each. Blots were developed with enhanced chemiluminescent (ECL) substrate (SuperSignal West Femto Maximum Sensitivity, Thermo Fisher) and visualized on the Odyssey UV imaging system (UVP). For matrix stiffness experiments, intracellular QSOX1 was normalized to GAPDH and secreted QSOX1 was normalized to ALG-2-interacting protein X (Alix). For hypoxic experiments, QSOX1 was internally normalized to β-tubulin and then normalized to normoxic controls.

### Immunofluorescence

PANC-1 cells were cultured on 22×22 mm coverslips (Electron Microscopy Sciences, no. 1.5) for 48 h and fixed with ice cold 4% PFA for 20 minutes. Samples were incubated in blocking buffer (1% BSA, 0.2% gelatin and 0.1% Tween-20) for 1 h at RT and then overnight at 4°C in primary antibody diluted in blocking buffer: anti-QSOX1 (1:1000, ProteinTech, 12713-1-AP), anti-HIF1α (1:500, 610958, (BD Biosciences)). Samples were washed and incubated with anti-rabbit and/or anti-mouse secondary antibody (1:1000; Thermo Fisher) and 647-phalloidin (1:160; Thermo Fisher) for 1 h at RT. Samples were counterstained with DAPI, mounted in Gelvatol, and visualized on a Zeiss LSM800 confocal microscope or a Zeiss Axio Observer Z1 epifluorescence microscope.

### Statistical analysis

Data is presented as mean ± SD. All statistical analysis was performed in GraphPad Prism 9 with α=0.05. For the ELISA, a 2-way ANOVA was performed with Šidák’s multiple comparisons. For the supernatant analyses, a Brown-Forsythe and Welch ANOVA was performed with Dunnett’s T3 multiple comparisons. For the RT-qPCR experiments, an RM one-way ANOVA with Geisser-Greenhouse correction and Tukey’s multiple comparisons test was performed. A p-value of P < 0.05 was considered statistically significant.

## RESULTS

### PAA gel formulations can successfully modulate cell attachment and morphology

To compare the moduli of our PAA gels with those reported in literature (see **Supplemental Table 1**), we polymerized our gels directly on a DHR3 rheometer using a PDMS ring that was positioned around the upper parallel plate. The PDMS ring minimized the introduction of oxygen, which is known to inhibit acrylamide polymerization by sequestering free radical generation. The storage modulus was recorded during until gels were fully polymerized (**Figure 2A**). The elastic moduli (E) were calculated from the storage modulus: gel 1 (2.32 kPa±0.28), gel 2 (42.8 kPa±0.97), and gel 3 (63.3 kPa±3.39). Gel 3 was consistently much lower than the reported value of 112.25 kPa ± 8.03. For ease of description for cell experiments, we labeled the gels as 2, 40, and 60 kPa (**Figure 2B**) throughout this paper.

**Figure 2:**
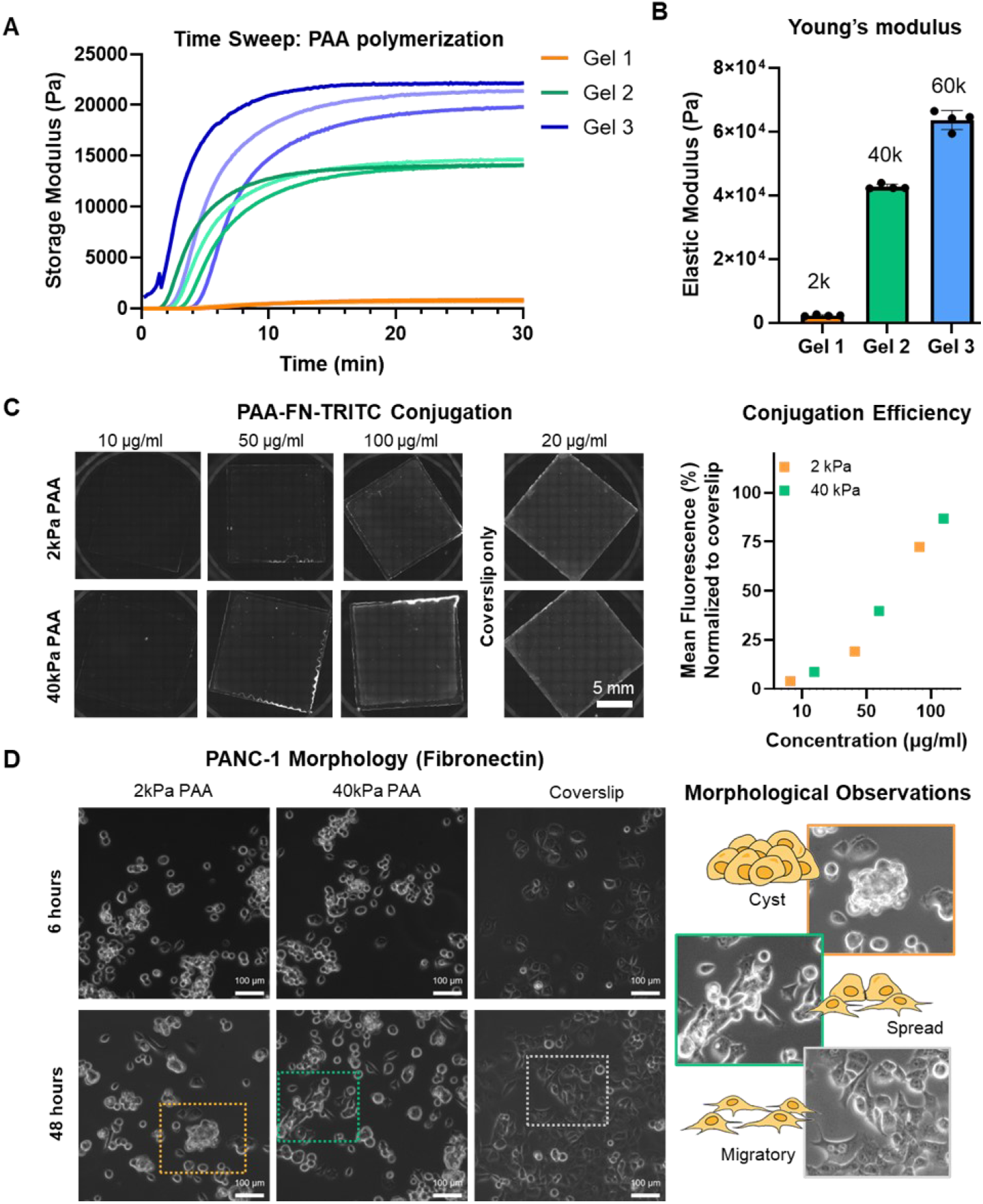
Characterization of PAA gels. (A) The storage modulus of PAA was measured with a rheometer using a time sweep with constant strain (0.1%) and oscillation (1 rad/s). (B) The elastic modulus was calculated for each gel (n=4). (C) ECM conjugation efficacy was visualized on gels using FN-TRITC and quantified for mean fluorescence intensity over the background. Signal was normalized to coverslips coated with 20 μg/ml FN-TRITC. (D) Brightfield images were taken 6 h and 48 h after initial seeding to assess cell morphology. The main morphological features were identified (cyst or clumped, spread, and migratory)

To confirm ECM attachment to the gels was uniform and complete, we conjugated FN-TRITC to PAA coverslips and imaged the entire region using an epifluorescence microscope with a 5x objective (**Figure 2C**). The mean fluorescence of the 2 kPa gel was slightly lower than 40 kPa gel for each concentration of FN-TRITC. This difference in fluorescence may be attributed to the lower concentration of acrylamide in the softer formulation, resulting in fewer available amide functional groups for covalent modification. At 100 μg/ml TRITC-FN, the PAA gels were at roughly 80% of the fluorescent intensity of the glass coverslips and were thus conjugated with ECM proteins at that concentration.

PANC-1 cells were seeded on PAA coverslips conjugated with fibronectin and collagen and imaged under brightfield to confirm differences in morphology (**Figure 2D, Supplemental Figure S1**). After 6 h, the cells on the PAA coverslips were attached, but had not yet started to spread. After 48 h, some cells on the PAA coverslips adopted a more spread morphology while others developed cysts. The 2kPa PAA gels were more likely to contain rounded or clumped cells. In contrast, the cells on the glass coverslips were already spread after 6 h and remained this way after 48 h.

### Intracellular QSOX1 levels are downregulated on soft PAA gels compared to cover glass

To visualize the cytoskeleton, PANC-1 cells were fixed after 48 h on fibronectin-conjugated coverslips and immunostained (**Figure 3A**). On the 2 kPa PAA coverslips, cells displayed an immature cytoskeleton and had little to no focal adhesions. On the 40 kPa PAA coverslips, cells had focal adhesions and local stress fibers, but displayed few cytoskeletal features. On the glass coverslips, cells displayed an organized cytoskeleton and contained mature focal adhesions. The glass coverslips also had abundant extracellular QSOX1 signal while the PAA coverslips did not. QSOX1 localized perinuclearly in all three conditions.

**Figure 3:**
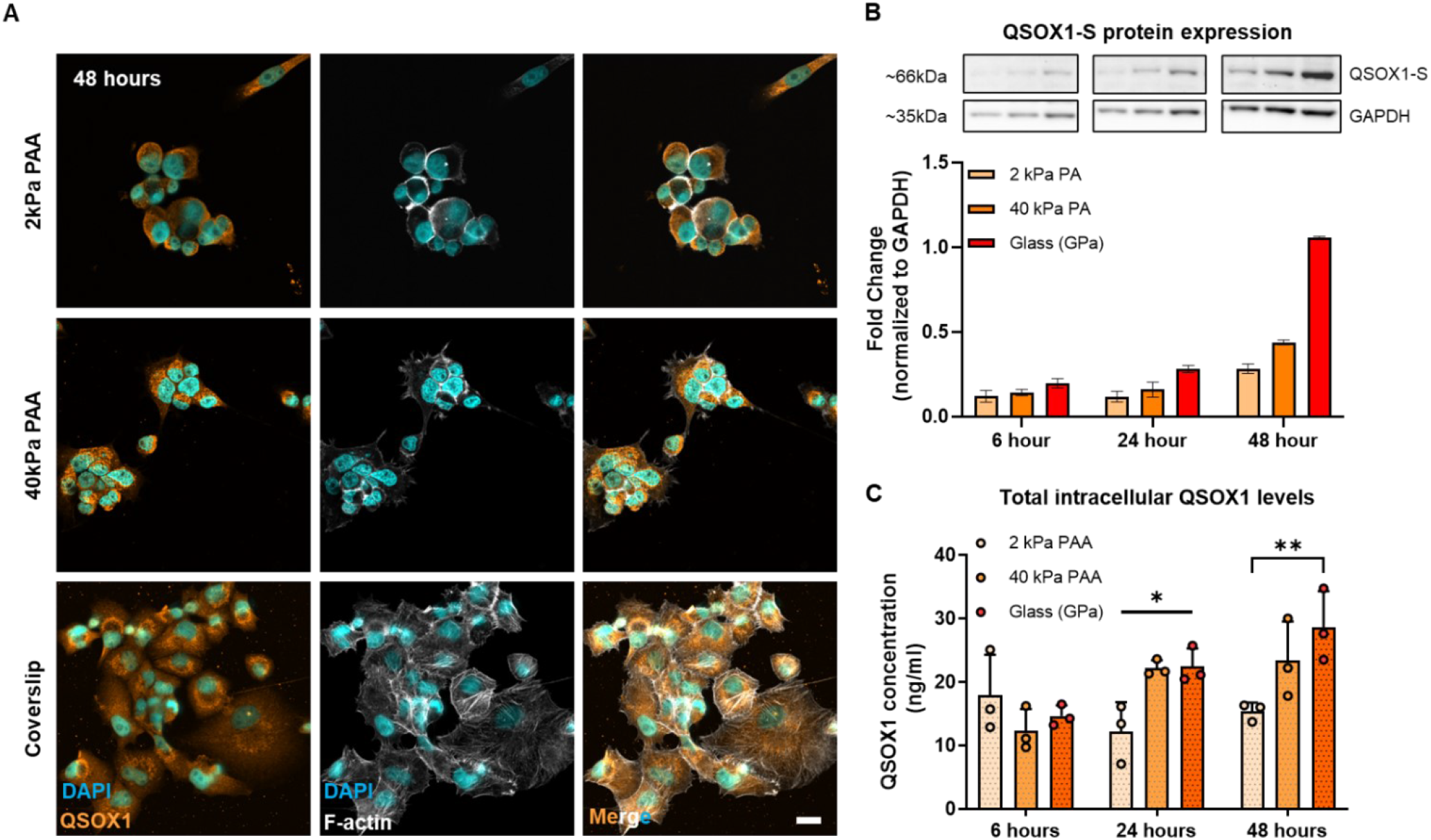
Soft surfaces downregulated intracellular QSOX1 concentration. (A) Cells on PAA gels and coverslips were immunostained for QSOX1 and visualized using confocal microscopy. Scale bar = 20 μm. (B) Western blot was performed, and QSOX1-S band intensity was quantified using densitometry. (C) Total intracellular QSOX1 concentrations were quantified using ELISA. Performed in experimental triplicate. *p<0.05 and **p<0.01

Western blot revealed that the cells cultured on fibronectin-coated glass coverslips had dramatically more intracellular QSOX1-S than the cells cultured on fibronectin-PAA coverslips after 48 hours (**Figure 3B**). This trend was also observed for cells cultured on gelatin-PAA coverslips (**Supplemental Figure S2**). Notably, total integrin β1 and QSOX1-L levels remained unchanged regardless of stiffness or ECM protein type. To quantify total intracellular QSOX1 concentrations, an indirect ELISA was designed in-house after extensive antibody validation and sample calibration (**Supplemental Figure S3**). ELISA revealed that intracellular QSOX1 levels were significantly higher on glass coverslips than on 2 kPa PAA coverslips after 24 h (p < 0.05) and 48 h (p < 0.01) (**Figure 3C**).

### QSOX1 secretion is decreased on PAA gels compared to ECM-coated dishes

To investigate whether matrix stiffness affected QSOX1 secretion from cells, conditioned media was collected after 24 h and the contents of the supernatant was analyzed^45^. The number of particles released from cells ranged from 200 to 500 particles/cell and was not statistically significant across all conditions (**Figure 4A**). The particle means likewise stayed consistent across all conditions (**Figure 4B**). The amount of protein released from cells was roughly 4 ng/cell for the ECM coated dishes and roughly 6 ng/cell for the 40 kPa and 60 kPa PAA dishes (**Figure 4C**). The amount of protein from the 2 kPa gel was extremely high (~38 ng/cell). The protein/particle ratio was calculated to assess relative biofouling of the samples (**Figure 4D**). The protein/particle ratio for ECM-coated dishes was roughly 13 fg/particle. The 40 kPa and 60 kPa PAA dishes had an average protein/particle of 30 fg/particle. The 2 kPa PAA dishes had an average of 90 fg/particle, suggesting significant exogenous protein contamination from FBS. Exogenous FBS contamination was confirmed after using a vehicle control (2 kPa PAA dish) that underwent the same procedure but contained no cells (**Supplemental Figure S4A**). However, QSOX1-S was not present in the vehicle control, excluding the potential interference from exogenous QSOX1. Additionally, NTA measured very few particles from FBS and the 2 kPa vehicle control compared to conditioned media generated from cells cultured on the 2 kPa gel (**Supplemental Figure S4B**), excluding potential exogenous particle contamination.

**Figure 4:**
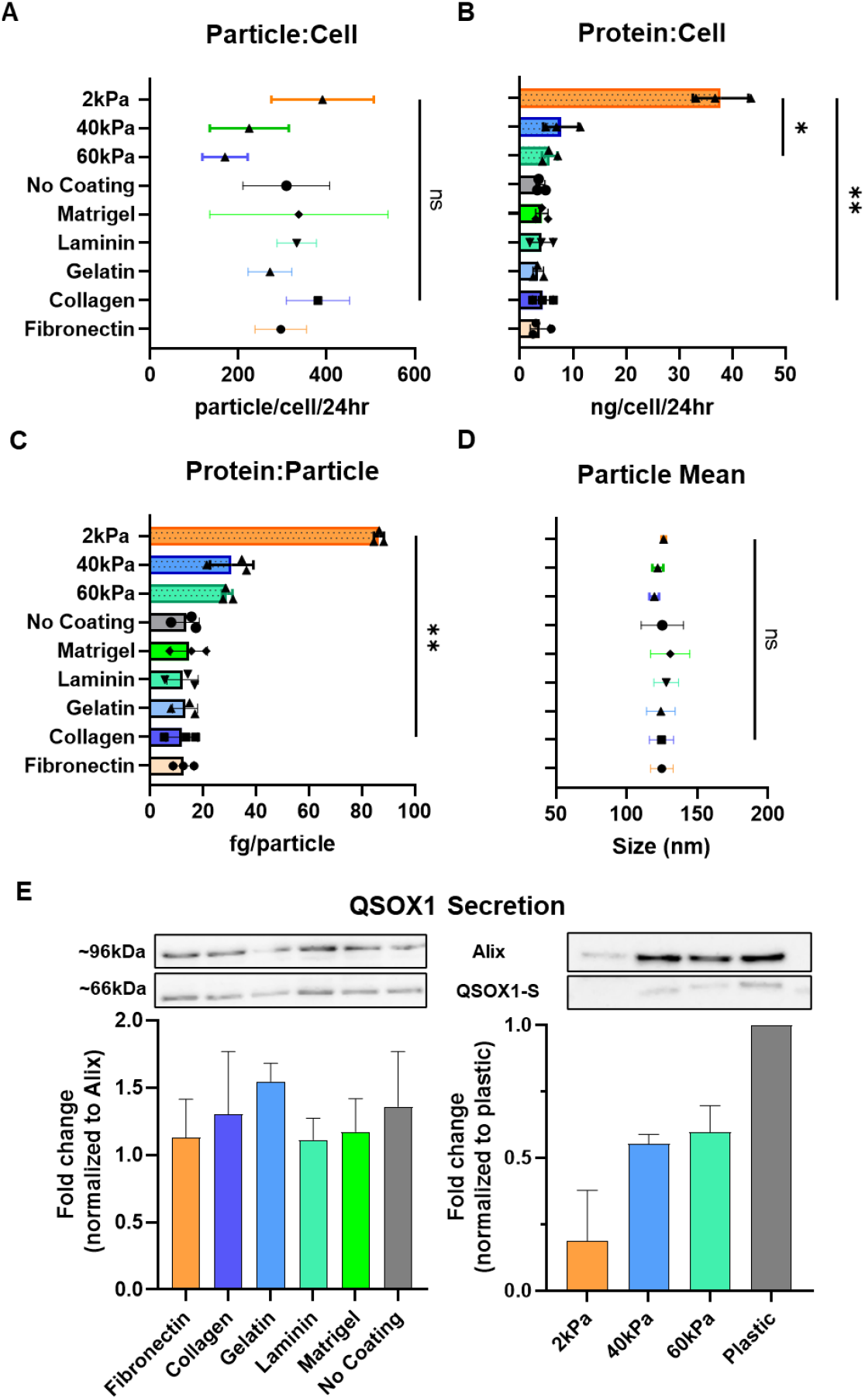
Analysis of conditioned media supernatant and secreted QSOX1. (A) Particle production: average number of particles released per cell over 24 h. (B) Protein secretion: average amount of protein secreted per cell over 24 h. (C) Protein/particle ratio indicating relative purity of samples. (D) Mean particle diameter. (E) Representative western blots and semi-quantification of QSOX1 relative to changes to the control protein Alix (n=2). *p<0.05 and **p<0.01. ns = not significant.

Since particle count was similar across all conditions, the particle-associated protein Alix was used as an internal control (**Figure 4E**). On ECM-coated dishes, QSOX1 accumulation in the supernatant was similar across all coatings. On PAA gels, there was less overall secreted QSOX1 in the supernatant of cells cultured on PAA gels compared to those on plastic surfaces.

### Hypoxic conditions have a limited effect on QSOX1 gene and protein expression over time

To determine whether hypoxia influenced QSOX1 mRNA levels, we performed RT-RTqPCR from cells that were either dosed with 400 μM CoCl_2_ or exposed to 1%O_2_ for up to 48 h (**Figure 5A**). Primers for QSOX1 and RPLP0 were tested and found to have efficiencies of 101% and 110% respectively (**Supplemental Figure S5**). For the first 24 h, we saw no differences in QSOX1 gene expression compared to time-matched controls. Even after 48 h, there was no significant increase in QSOX1 levels for either type of hypoxic induction. Interestingly, QSOX1 gene expression varied over time more than it varied in response to hypoxia. When each replicate was normalized to the baseline time measurement, QSOX1 gene expression changed slightly depending on the experimental replicate but followed a similar trend between hypoxic and normoxic cells for each run (**Figure 5B)**.

**Figure 5:**
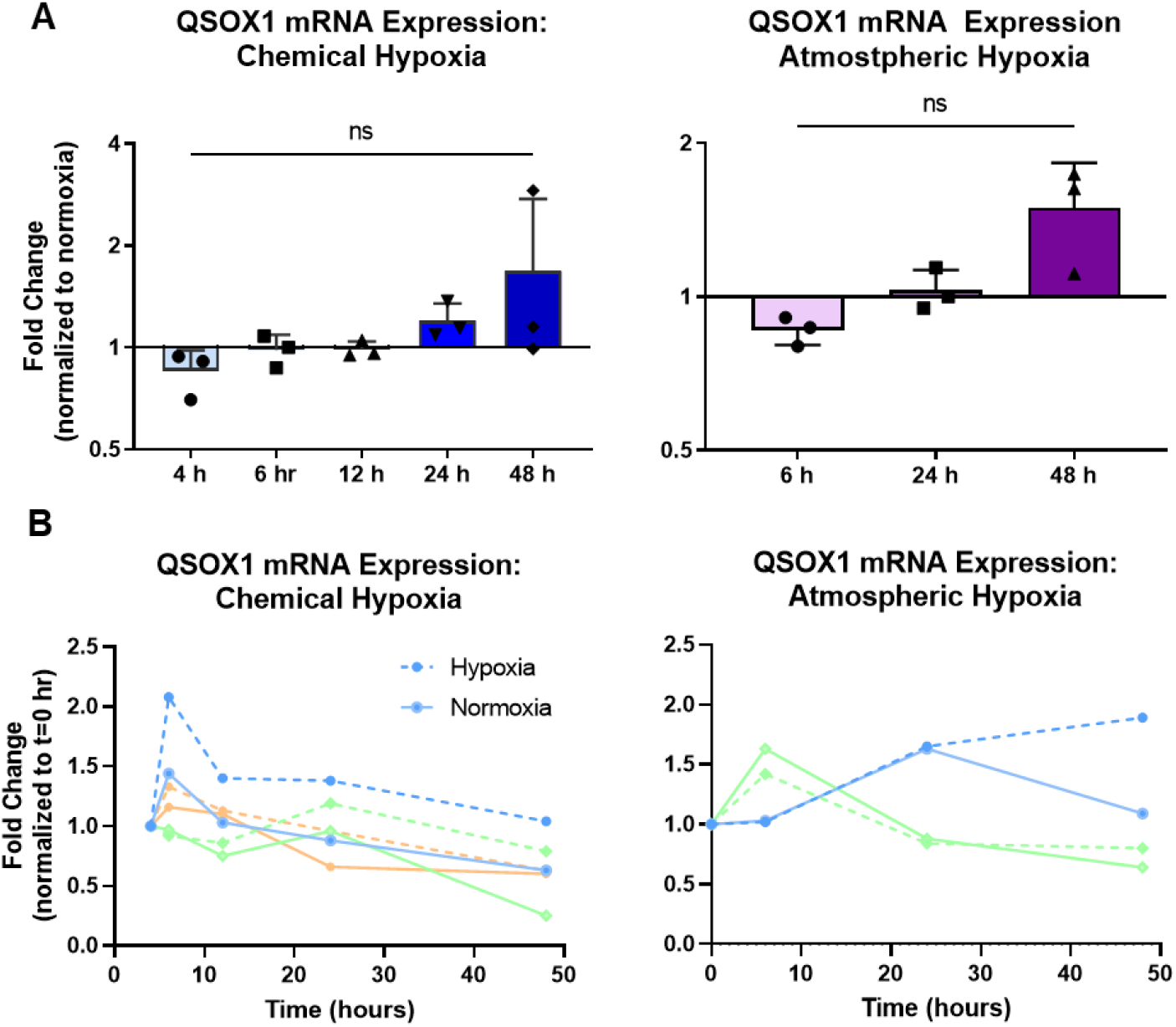
Temporal changes to QSOX1 gene expression under hypoxic conditions. (A) Fold change of QSOX1 gene expression normalized to time-matched normoxic controls for chemical and atmospheric induction of hypoxia (n=3). ns = not significant. (B) Fold change in QSOX1 gene expression normalized to t = 0 h for normoxic and hypoxic conditions.

We quantified intracellular QSOX1 protein expression after chemical induction of hypoxia (**Figure 6A**). We found no clear changes in either QSOX1-L or QSOX1-S levels over 48 h. To confirm that we were stimulating a hypoxic response from cells, we also semi-quantified changes to HIF-1α protein levels compared to time-matched controls. There was a dramatic increase in HIF-1α that peaked at 6 h for both replicates. We fixed and immunostained cells after 48 h to determine intracellular localization and saw a clear nuclear translocation of HIF-1α in the hypoxic conditions (**Figure 6B**). QSOX1 remained perinuclear for both conditions.

**Figure 6:**
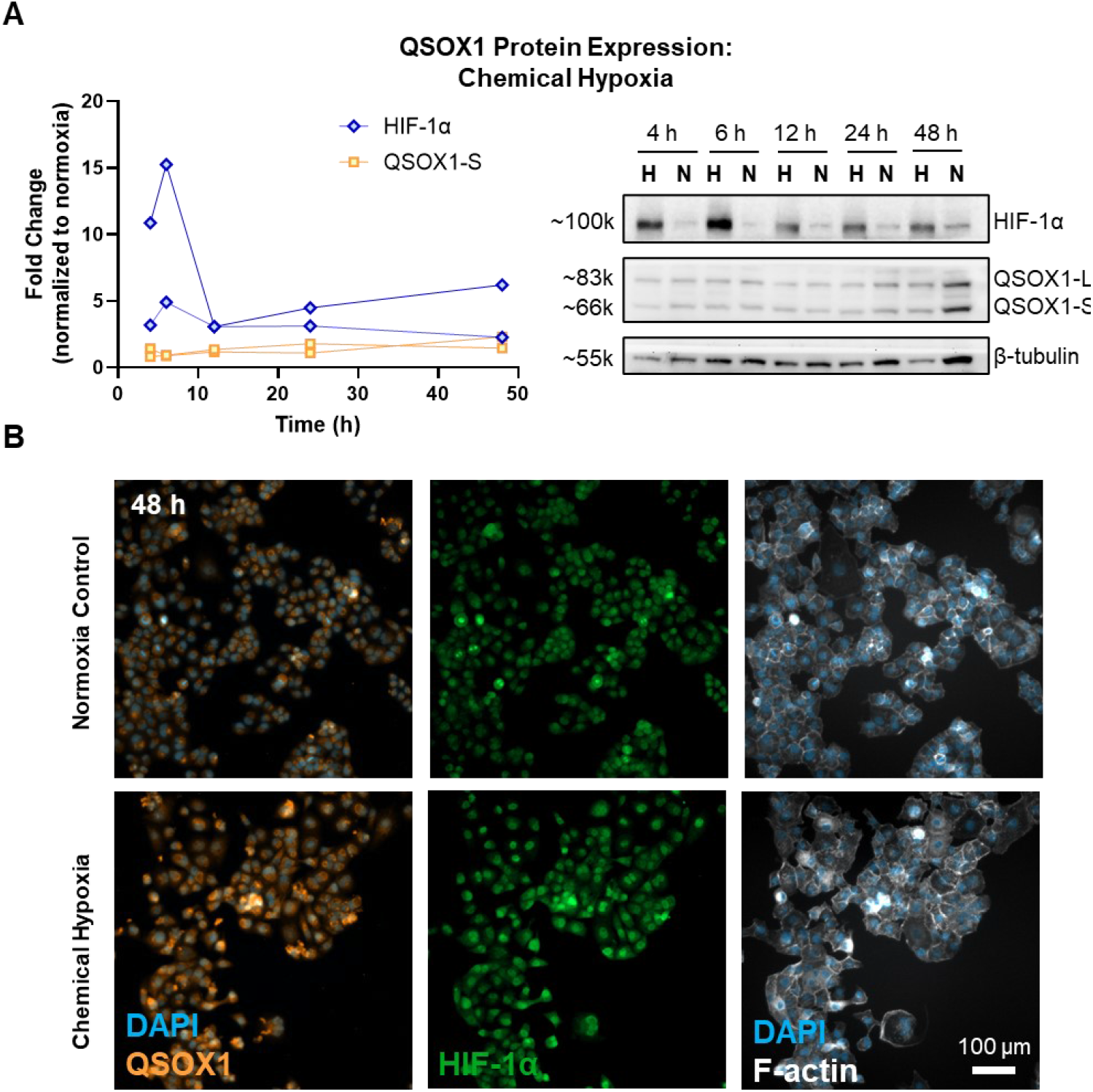
Temporal changes to intracellular QSOX1 and HIF-1α protein levels under hypoxia. (A) Fold change of intracellular QSOX1 and HIF-1α protein normalized to time-matched normoxic controls (n=2). Representative western blot (H=hypoxia, N=normoxia). (B) Cells under hypoxic and normoxic conditions were fixed and immunostained for QSOX1 and HIF-1α and visualized with an epifluorescence light microscope under a 20x objective.

## DISCUSSION

The key feature of PDAC is the presence of a strong desmoplastic reaction, or fibrotic response, of the stroma of pancreatic lesions (**Figure 7A**). The stroma is estimated to comprise more than 80% of the tumor mass and contains a heterogenous population of cells and extracellular components^50,51^. The deposition of dense ECM protein in the stroma results in a mechanical stiffening of the tumor, poor vascularization and nutrient perfusion, and severe hypoxia^5^. Over 90% of all PDAC diagnoses are regionally or distantly metastatic^52^, which complicates treatment options. In the 2000s, therapeutics and adjuvants were developed to target and deplete the tumor stroma to relieve pressure, decrease matrix density, and increase vascular perfusion. It was presumed that ablating the stroma would enhance chemotherapy drug performance and therefore increase survival.^16,53,54^. Many of these adjuvant therapeutics showed promising results in pre-clinical trials, only to fail in the clinic^55,56^. The only treatment that has increased median survival of patients with metastatic PDAC is FOLFIRINOX (folinic acid, fluorouracil, irinotecan, and oxaliplatin); however, this combination of chemotherapy agents is only indicated for patients with good performance due to its serious side effects^57,58^. To date, no adjuvant therapies have substantially increased the median survival of patients with metastatic PDAC.

**Figure 7:**
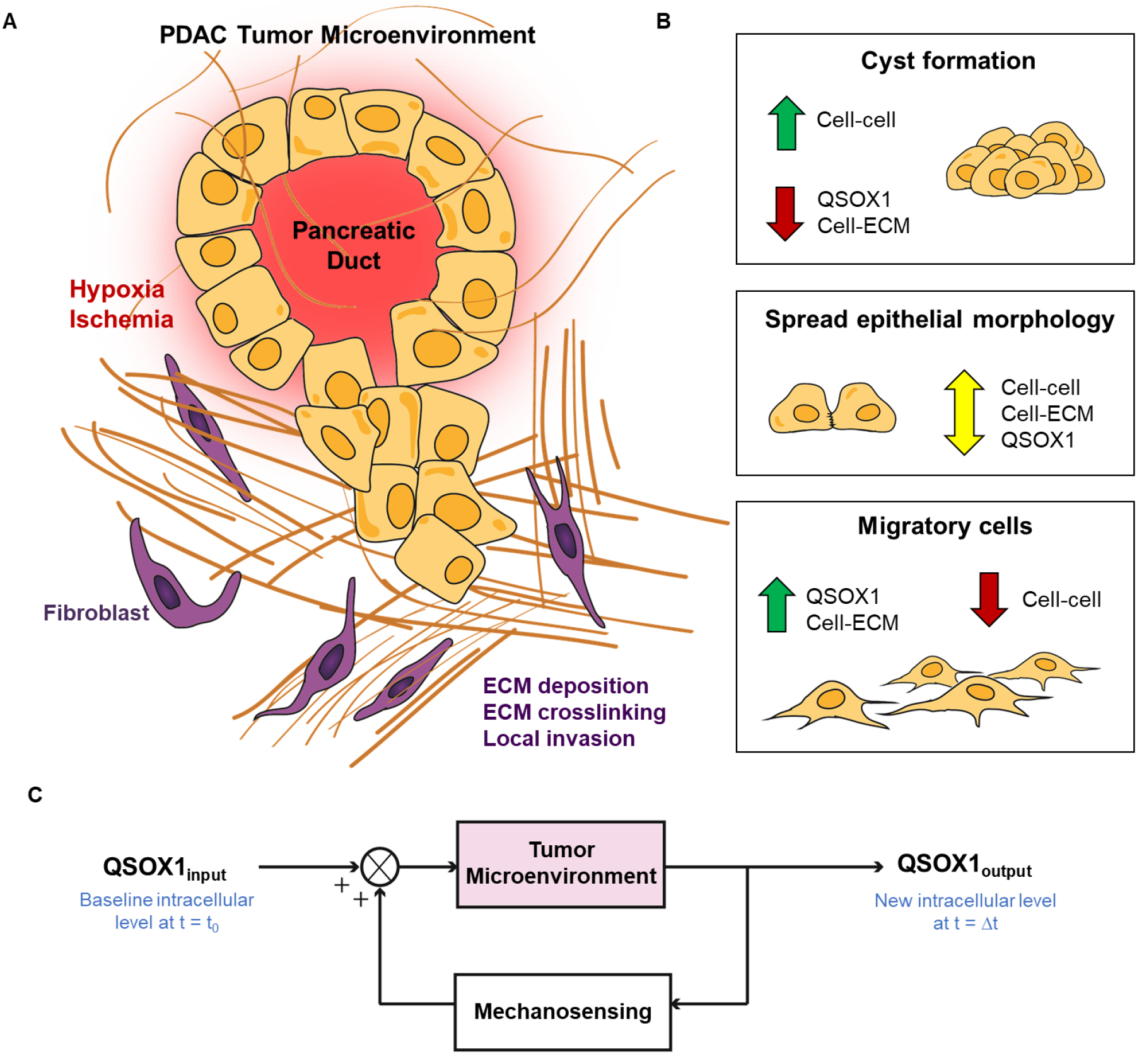
Proposed effect of the tumor microenvironment on QSOX1 regulation. (A) PDAC is characterized by maladaptive changes to the pancreatic tissue, which results in hypoxia, ischemia, and altered matrix mechanics. Cells undergo behavioral changes in response to these challenges, which are visualized by their morphology and phenotype. (B) There were three main cell morphologies observed on the PAA gels and glass coverslips: cystic or clumped, spread, and migratory. It is proposed that the more spread and migratory cell phenotype is correlated with the upregulation of QSOX1. Cells that favored cell-cell contacts over cell-ECM contacts on soft surfaces downregulated QSOX1. (C) A proposed diagram representing a feedback system in which cells increase QSOX1 levels based on the mechanical features of the tumor microenvironment. At an initial time, t=0, pancreatic cancer cells have baseline QSOX1 levels. When the extracellular environment changes, cells can sense the changes to the matrix (e.g., increased tensile strength or altered ECM protein deposition) and react by increasing intracellular QSOX1 levels. QSOX1 levels do not increase when cells are on softer substrates with limited cell-ECM contacts, and thus, these levels remain at baseline concentrations.

New efforts have been made to tease apart individual contributors to the tumor microenvironment with the goal of exerting a more precise manipulation of the stroma^59^. Matrix remodeling enzymes have emerged as potential therapeutic targets, including QSOX1. The first connection between QSOX1 and PDAC was made in 2009 when a peptide fragment corresponding to QSOX1-L was found in the serum of patients^60^. Molecular inhibitors of QSOX1 have since shown anti-tumor activity in mice, including decreased tumor growth and local invasion^24,27,30^. In the span of less than a decade, QSOX1 went from a relatively obscure enzyme to the center of attention in dozens of cancer-related studies. Despite this recent surge of publications, it is still unclear why QSOX1 is overexpressed in solid tumors and its biological functions outside the cell, although there are several related functions that have been described.

One interesting observation is that QSOX1 seems to be extrinsically linked to cell-ECM interactions. When QSOX1 was silenced in lung fibroblasts, the fibroblasts detached from monolayers and floated, but they were viable and could be reattached with the addition of exogenous QSOX1^31^. Inhibiting QSOX1 also depleted laminin in confluent monolayers, increased the presence solubilized laminin, and decreased invasive potential in lung cancer cells. In another study, silencing QSOX1 in pancreatic tumor cells resulted in decreased invasion and a corresponding inactivation of MMP-2 and MMP-9^23^. Most studies employ the standard protocol of inhibiting QSOX1 (antibody, molecular, or RNA) to observe subsequent changes in cell behavior^25,29,61^. While QSOX1 has been manipulated to observe changes in cell-ECM interactions, the matrix has not been manipulated to observe changes in QSOX1 expression. Our data would suggest that the tumor microenvironment itself can modulate QSOX1.

When we cultured PANC-1 cells on PAA gels versus glass coverslips, intracellular QSOX1-S concentrations significantly increased on the coverslips. Interestingly, we observed that total intracellular QSOX1 protein concentrations started at a baseline level on all surfaces 6 h post-seeding. After 24 h, QSOX1 concentration on the 40 kPa PAA gel and coverslip increased. Finally, after 48 h, QSOX1 on the coverslip increased to about double its initial concentration. The 2 kPa gels never saw an increase in QSOX1 from the initial timepoint. We speculate that the change in QSOX1 concentration was a function of cell spreading and/or mobility (**Figure 7B**). After 48 h, cells on the coverslips had spread and formed mature, organized actin filaments and focal adhesions. Conversely, cells on the 2 kPa gel had few focal adhesions, no clear actin structure, adopted a rounded phenotype, and tended to form cysts. Cells on the 40 kPa gels were between these two phenotypes. Given the connection between QSOX1 expression and cell adhesion and migratory behavior, it is possible that stiff surfaces, like plastic or glass, upregulate QSOX1 production while soft surfaces, like PAA, do not. Such a feedback system could explain why QSOX1 is overexpressed in high grade solid tumors (**Figure 7C**)^26,29,62^. We propose that the ability for cells to move, spread, and interact with the ECM has a significant effect on QSOX1-S regulation. We did not see any discernable changes to QSOX1-L, the non-secreted isoform containing a transmembrane domain.

Because intracellular QSOX1-S levels were downregulated on softer substrates, we next investigated if this translated to less secreted QSOX1-S. There were some major limitations inherent to our large PAA dish platform. First and foremost, there was significant exogenous serum protein fouling in the 2 kPa PAA gels despite the use of basal medium during conditioned media production. Cells were initially cultured on PAA gels for 24 h in DMEM-ITS media (containing 2% FBS) to facilitate attachment and viability. Care was taken to minimize biofouling. We attempted to remove the serum proteins that diffused into the PAA gels by soaking them in HBSS++ for 2 hours before adding basal DMEM, which was mostly successful in the 40k Pa (8% acrylamide) and 60 kPa (12% acrylamide) gels, but not in the 2 kPa (4% acrylamide) gels. We then included an acellular process control for the 2 kPa gel and found QSOX1-L contamination (from FBS), but no exogenous QSOX1-S in the supernatant. Because protein quantification was unreliable, we quantified particle secretion from cells in all conditions. We found no statistical differences, and thus used Alix as an internal protein control for our western blots and loaded samples based on particle count. Overall, we found a decrease in QSOX1 accumulation in the supernatant from cells cultured on the 40 kPa and 60 kPa gels compared to tissue culture plastic, and we tentatively report the same trend for the 2 kPa gels. There is most likely a causal relationship between intracellular and secreted QSOX1 concentrations, but further investigation is warranted. One last limitation of our experimental design was the inability to quantify how much protein (specifically QSOX1) may have become trapped in the PAA gel after conditioned media collection. Coomassie stain showed little to no protein remaining in the gel after cells were trypsinized and removed for counting, but this observation was qualitative at best.

Finally, we simulated hypoxia in our PANC-1 cells to corroborate an earlier finding that QSOX1 expression is upregulated under hypoxic conditions in a HIF-1α-dependent manner^44^. Previously, two functional HREs were identified in the QSOX1 gene. Unfortunately, we could not replicate these findings. Instead, we found that QSOX1 gene and protein expression remained unchanged during the first 24 h with atmospheric and chemical induction of hypoxia. We quantified HIF-1α and noted an intense peak at 6 h followed by a plateau after 12 h, a characteristic response to hypoxia that has been well-documented^63,64^. There were no notable changes in QSOX1 protein levels. When we normalized QSOX1 gene expression to the baseline starting levels (t=0) rather than to time-matched controls, there was a strong temporal component to the fluctuations. We cannot reconcile these differences in the absence of primer sequences and the antibodies used in that study^44^. Primers from our experiments were tested and validated (**Supplemental Figure S5**) and our QSOX1 antibodies were characterized (**Supplemental Figure S6**) to support our conclusions.

In summary, we present novel findings that suggest QSOX1 is regulated by matrix mechanics, but not necessarily hypoxia. The desmoplastic response in the PDAC stroma leads to increased stiffness, hypoxia, and other maladaptive changes. However, there is complex interplay between signaling cascades involved in hypoxia, fibrosis, and matrix remodeling during tumor progression. Here, we decoupled two characteristics of the tumor microenvironment to determine which one predominantly influences QSOX1 expression and found that matrix mechanics directly modulates QSOX1. Understanding which signaling cues dominate QSOX1 can inform future clinical directions for targeting the tumor stroma as well as understanding the pathological progression of PDAC. QSOX1 is likely a significant matrix remodeling enzyme in malignant solid tumors and deserves continued attention.

## Abbreviations

PDAC: Pancreatic ductal adenocarcinoma
ECM: extracellular matrix
LOX: lysyl oxidase
QSOX1: quiescin sulfhydryl oxidase 1
MMP: matrix metalloproteinases
PAA: polyacrylamide
HIF-1: hypoxia inducible factor 1
PDMS: polydimethylsiloxane
APTES: 3-aminopropyltriethoxysilane
APS: ammonium persulfate
TEMED: tetramethylethylenediamine
sulfo-SANPAH: sulfosuccinimidyl 6-(4’-azido-2’-nitrophenylamino)hexanoate
TRITC: 5(6)-tetramethylrhodamine isothiocyanate
HBS: HEPES-buffered saline
NTA: nanoparticle tracking analysis
LDS: lithium dodecyl sulfate.

## ACKNOWLEDGMENTS

This work was funded by the National Institutes of Health (R01GM26643, U54GM104941). The authors would also like to acknowledge Bryan Ferrick for his assistance with rheological experiments. The rheometer is supported by the National Institute of General Medical Sciences (P30GM110758) through the Molecular Design of Advanced Biomaterials Center of Biomedical Research Excellence (COBRE) at the University of Delaware.

## CONFLICTS OF INTEREST

The authors have no conflicts of interest to disclose.

## AVAILABILITY OF SUPPORTING DATA

The supporting data and materials are provided in Supplementary Figures and Tables.

## AUTHOR CONTRIBUTIONS

CMH conceptualized the project, performed the experiments, analyzed data, and wrote the manuscript. CT contributed original observations and insight regarding QSOX1 catalysis. JPG conceptualized the project, supervised the experiments, and wrote the manuscript. All authors read, edited, and approved the final manuscript.

## Notes

### Competing Interest Statement

The authors have declared no competing interest.

