## Supplemental Figures and Tables for "Matrix mechanics, not hypoxia, modulate quiescin sulfhydryl oxidase 1 (QSOX1) in pancreatic tumor cells"

*bioRxiv*

Table 1: PAA formulations with their reported and measured elastic moduli.

| Acrylamide (%) | Bis-acrylamide (%) | Reported value<br>E ± SD (kPa) | Measured value<br>E ± SD (kPa) |
|---|---|---|---|
| 4 | 0.1 | 2.01 ± 0.75[1] | 2.32 kPa±0.28 |
| 8 | 0.48 | 40.4 ± 2.39[1] | 42.8 kPa±0.97 |
| 12 | 0.25 | 112.25 ± 8.03[2] | 63.3kPa±3.39 |

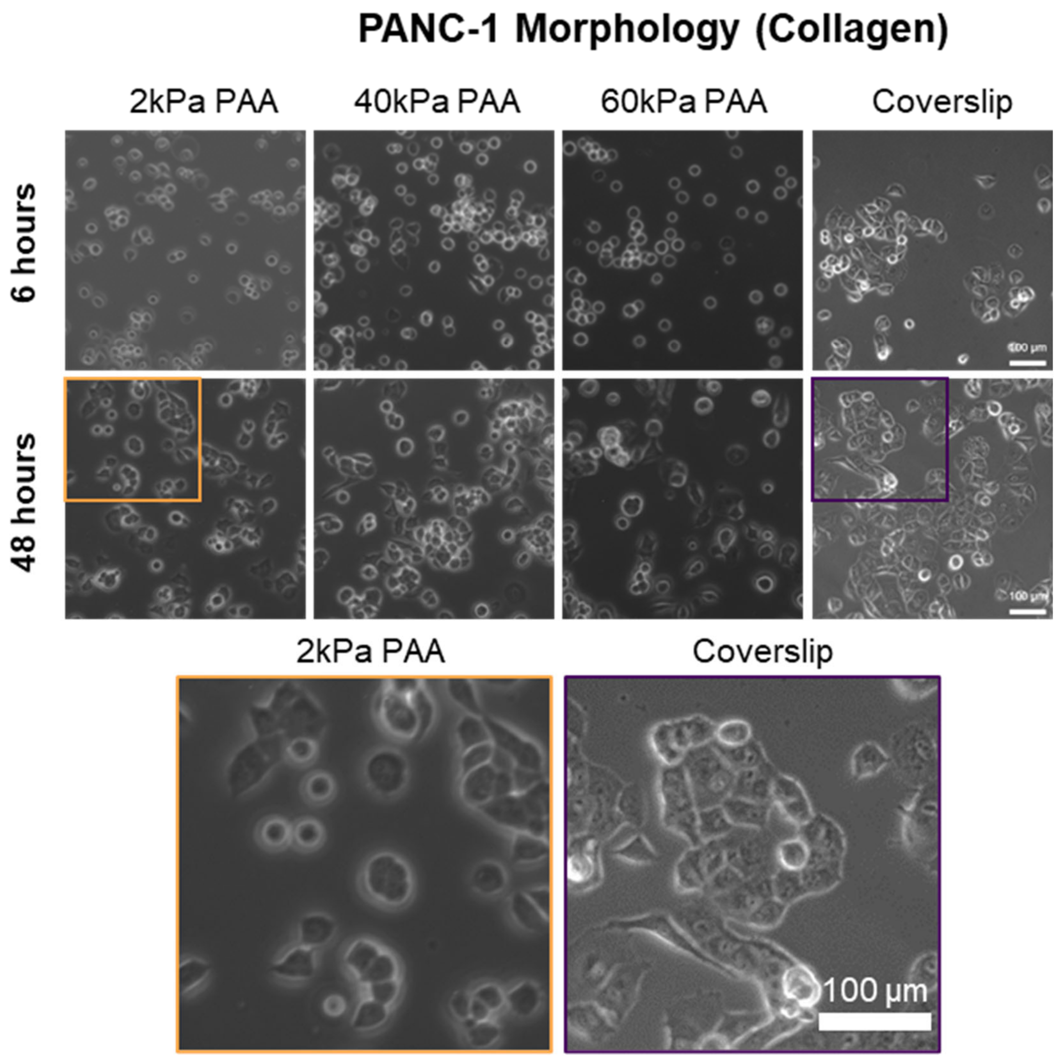


**Figure S1: PANC-1 morphology on collagen-PAA gels.** Brightfield images were taken 6 h and 48 h after initial seeding to assess cell morphology..

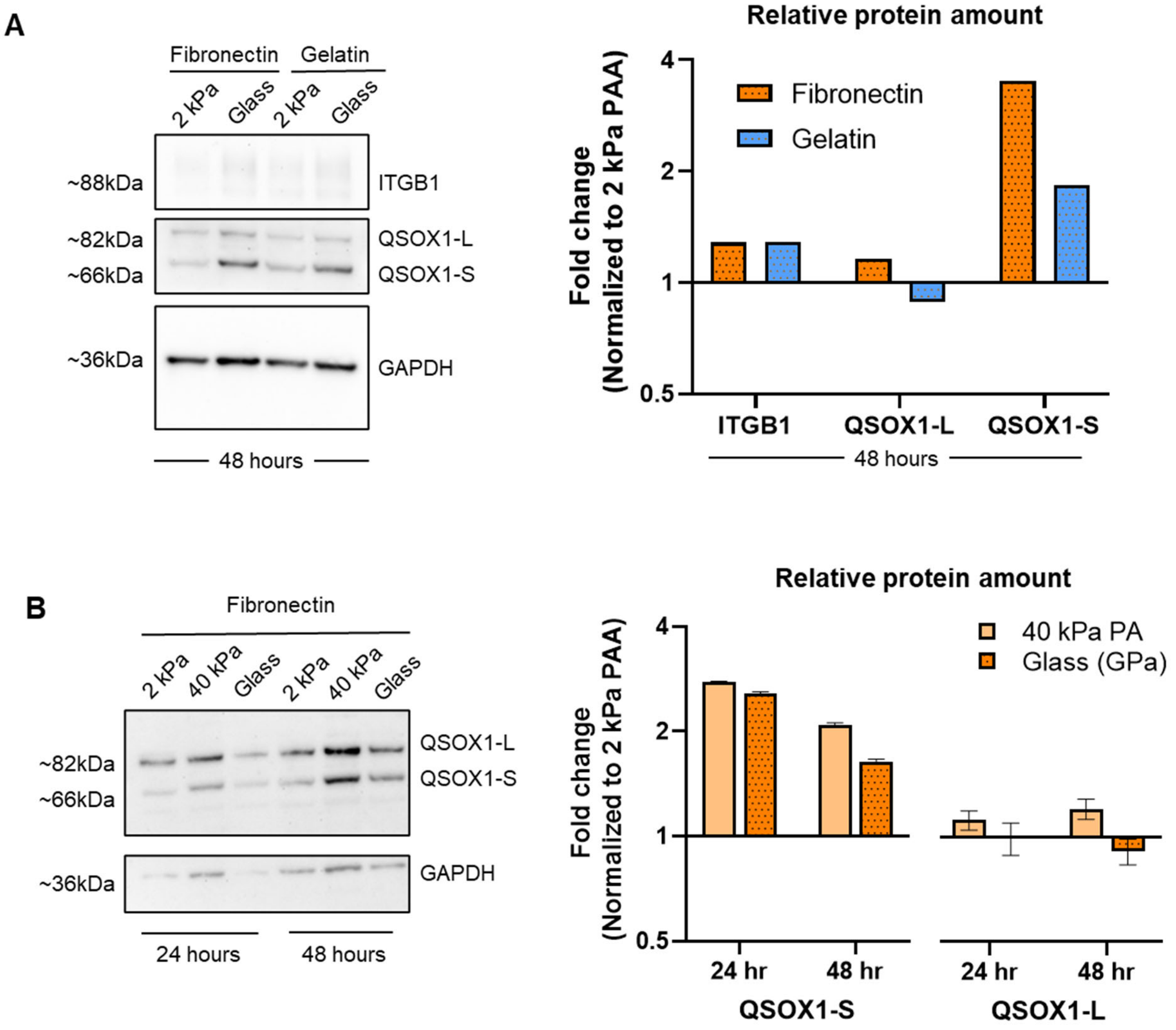


**Figure S2: Intracellular QSOX1-S levels are increased on glass coverslips compared to soft (2 kPa) PAA after 48 hours, but not QSOX1-L or ITGB1.** (a) Western blot of ITGB1, QSOX1, and GAPDH from cells cultured on fibronectin or gelatin coated coverslips or PAA gels for 48 hours. Measurements were internally normalized to GAPDH. QSOX1 and ITGB1 were experimentally normalized to the 2 kPa PAA condition. (b) Western blot of QSOX1 and GAPDH from cells cultured on fibronectin coated coverslips or PAA gels for up to 48 hours. Measurements were internally normalized to GAPDH. QSOX1 was experimentally normalized to the 2 kPa PAA condition to show relative changes to QSOX1-L and QSOX1-S.

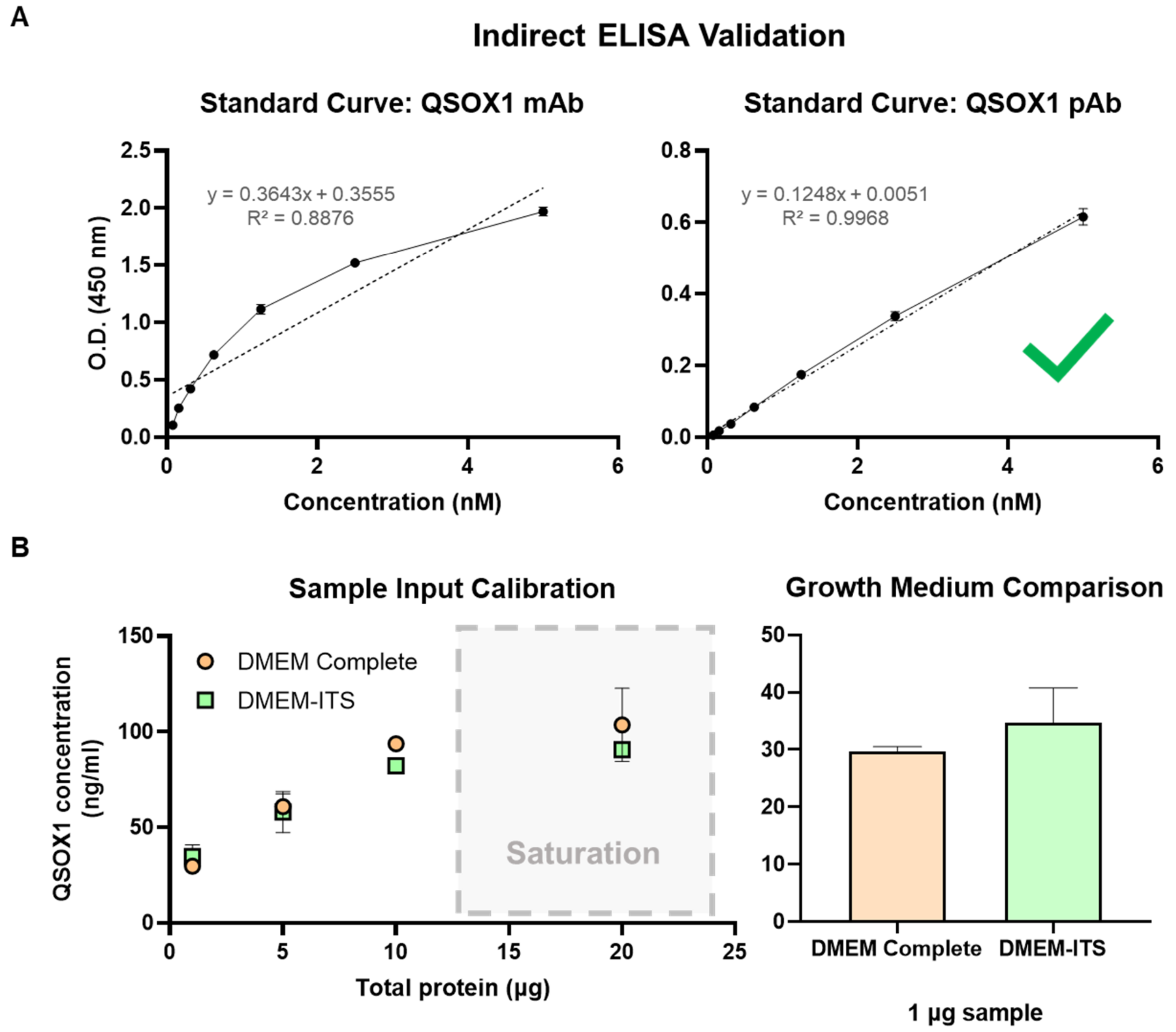


**Figure S3**: **Indirect ELISA validation.** (a) Standard curves were generated from two commercially available antibodies, QSOX1 mAb (Abcam) and QSOX1 pAb (ProteinTech). The QSOX1 pAb was selected based on linearity and sensitivity. (b) Serial dilutions of cell lysate were used to determine the range of sample input for the ELISA in a 96-well format and its saturation point and/or inhibitory effects. The 1 µg sample input was selected. There were no changes to baseline QSOX1 levels in the low serum PANC-1 cell line compared to the parental line.

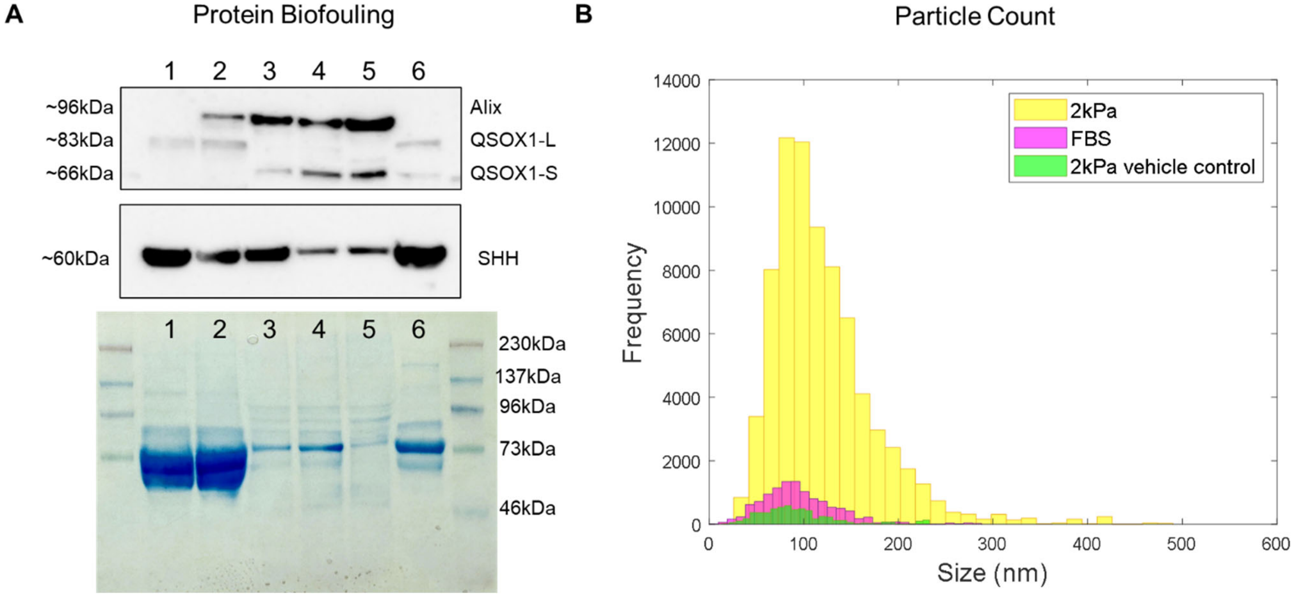


**Figure S4**: **Additional observations of protein and particle levels in conditioned media**. (a) Cells were cultured in DMEM-ITS (containing 2% FBS) and then grown in basal media for 24 hours. Conditioned media was collected and concentrated. Western blots of Alix, QSOX1, and SHH and the corresponding Coomassie stained gel showing total protein. Lanes were loaded based on particle count, not protein concentration; however, less sample from the 2 kPa PAA conditions had to be loaded on account of high exogenous protein amount (>3 mg/ml). The lanes are (1) 2kPa PAA (vehicle control), (2) 2 kPa PAA, (3) 40 kPa PAA, (4) 60 kPa PAA, (5) tissue culture plastic, and (6) FBS. The vehicle control underwent the same experimental process but contained no cells. (b) NTA of FBS, 2 kPa PAA, and the 2 kPa PAA control (vehicle control) showing particle size distributions and concentrations. The same concentrations of FBS and sample were used between western blot and NTA for direct comparison.

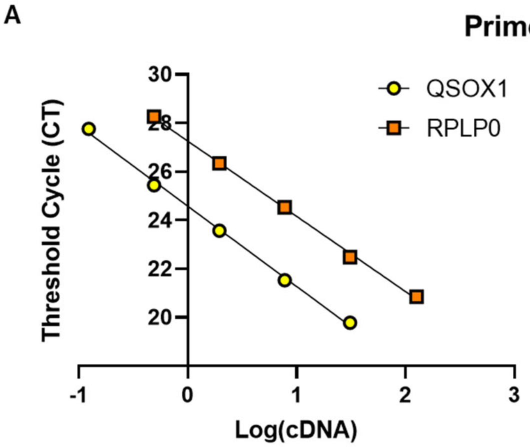


| | QSOX1 (Fam) | RPLP0 (Vic) |
|---|---|---|
| Equation: | Y = -3.307*X + 24.58 | Y = -3.098*X + 27.26 |
| $R^2$ | 0.9978 | 0.9989 |
| Efficiency: | 1.01 | 1.10 |

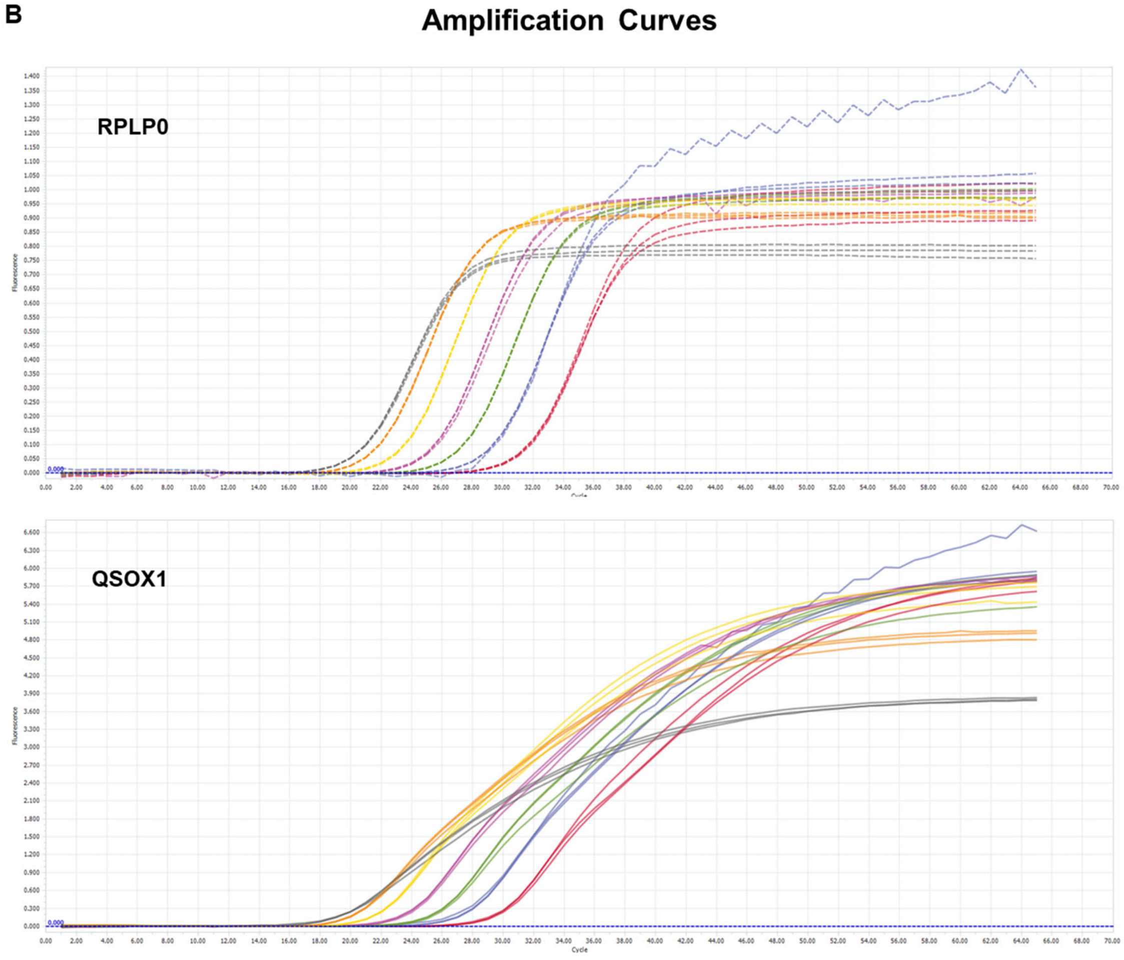


**Figure S5: Primer efficiency and probe performance**. (a) The $C_t$ values were plotted against the log of the template (cDNA) concentration and the efficiency was calculated based on the equation (Primer efficiency = $10^{(-1/slope)}$ -1). (b) Fluorescence versus cycle number for the reference (RPLP0) and target (QSOX1) genes using a serial dilution of the template (mRNA isolated from PANC-1 cells).

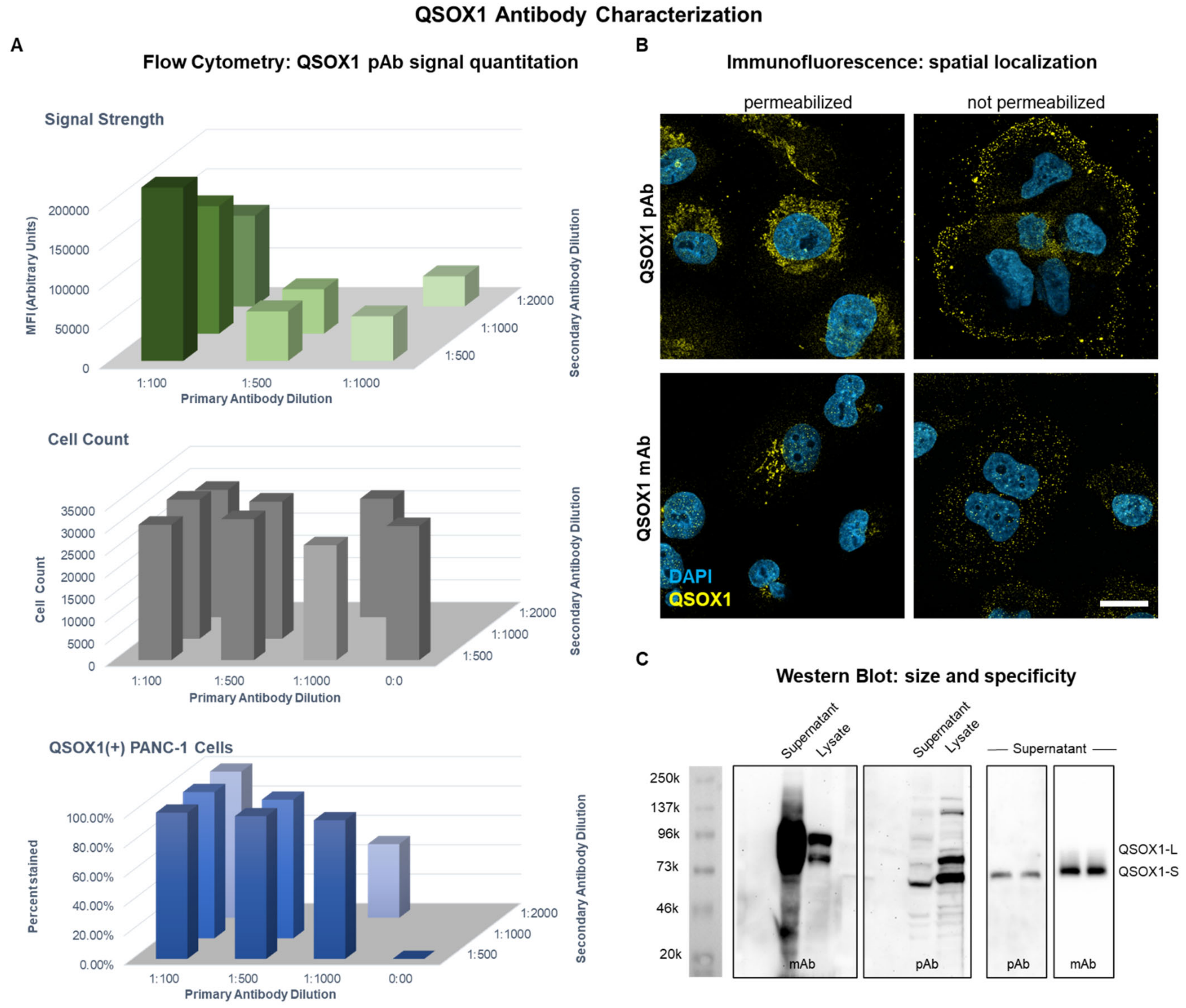


**Figure S6: QSOX1 antibody validation.** (a) PANC-1 cells were grown in a 24-well plate, trypsinized, fixed, permeabilized with 0.1% saponin, and immunostained with QSOX1 pAb antibody (ProteinTech) and 488-conjugated secondary with the indicated dilutions. Fluorescent intensity, cell number, and percent of positively stained cells were quantified using flow cytometry (n=3 technical replicates). (b) Cells grown on glass coverslips were fixed with 4% PFA for 15 min and were either permeabilized with Triton X-100 or directly immunostained with anti-QSOX1 mAb or pAb. Cells were counterstained with DAPI and visualized using confocal microscopy with a 63x oil objective. Scale bar = 20 µm. (c) Cell lysate or supernatant (from concentrated conditioned media) was immunoblotted with anti-QSOX1 mAb or pAb.
